# Using camouflage for conservation: colour change in juvenile European lobster

**DOI:** 10.1101/431692

**Authors:** Sara Mynott, Carly Daniels, Stephen Widdicombe, Martin Stevens

**Author notes:** Corresponding author: (SM).

## Abstract

Changes in coloration enable animals to refine their camouflage to match different visual environments. Such plasticity provides ecological benefits and could potentially be exploited to support conservation or stock enhancement efforts. One application could be ensuring that hatchery-reared animals, reared to stock wild populations, are appropriately matched to their environment on release. European lobster (*Homarus gammarus*) hatcheries aim to restock or enhance local lobster populations by rearing juveniles through their most vulnerable stages, then releasing them into the wild. However, little consideration has been given to their camouflage and the implications of matching individuals to their release site. This study assesses to what extent juvenile lobsters can change coloration to match their background and whether hatchery practices could be altered to enhance lobster camouflage. We test this by switching individuals between black or white backgrounds in the laboratory and monitoring their coloration over time. Our work demonstrates the capacity of juvenile lobsters to change lightness in response to their surroundings. We show that juvenile lobsters are capable of small changes in luminance (perceived lightness) to better match their background over 2-3 weeks. These changes potentially correspond to improved camouflage, based on a model of predator (European pollack, *Pollachius pollachius*) vision. However, over a longer period (5 weeks), lobsters maintained on either background converged on the same darker coloration, suggesting that lobsters also experience changes in appearance associated with ontogeny. By refining the approaches used here, there is potential for hatcheries to rear lobsters on backgrounds that better match their release site. However, such manipulations should be considered in the context of ontogenetic changes and release timing (which varies between stocking programmes). This study highlights the potential to use colour change in stocking and aquaculture, as well as gaps that could be addressed through further research in this area.

## Introduction

Background matching is one of the most widely used anti-predator defence strategies in nature (1), with many species using colours and patterns to match their surroundings and avoid detection and predation. In a wide range of taxa, both terrestrial and aquatic, camouflage can be achieved through plastic changes in appearance (2,3). This enables animals to respond to either fast and unpredictable changes in the visual environment through rapid (seconds and minutes) colour change, or slower and more predictable environmental change with gradual (hours, days, and weeks) appearance changes (4). One of the most widely studied groups, particularly in terms of the mechanisms and functions of colour change, are the crustaceans (3,5). Many crustacean species employ camouflage to conceal themselves from predators and several have been shown to change colour to better match their background, including crabs (6–8), prawns (9,10) and isopods (11,12), leading to phenotype-environment matches (13,14). It is likely that many juvenile crustaceans are phenotypically plastic with the ability to match the environment in which they settle (15,16). Such plasticity in background matching should confer a substantial survival advantage (17,18), and is likely to be particularly important for species and life stages that otherwise have limited anti-predator defences. The benefits of understanding camouflage and how it works have long been realised in applied areas such as the military and art and design (19,20), and colour change applications are growing in other fields, such as biomimicry (21) and animal welfare (22). However, an important application, seldom considered, is how an understanding of camouflage and colour change may be harnessed for conservation. For example, how colour change for camouflage could be applied in captive breeding and stocking programmes to improve post release survival and therefore stocking success.

Stocking, including stock enhancement and restocking, is used around the world to meet conservation needs and future seafood demands (23–25). Such programs improve and sustain capture fisheries *via* the release of cultured juveniles into the wild. There are some 180 cultivated marine species worldwide (24). These are reared in captivity (when they are most vulnerable to predation) before release into the wild (at a larger, more resilient size) in order to overcome challenges in recruitment and restore spawning stock biomass (26). While extensive research has been carried out to ensure the viability of released individuals, little consideration has been given to their ecology on release (27), in particular to the development of appropriate anti-predator defence behaviours. Unlike their wild counterparts, hatchery-reared juveniles are naïve to predators, which often puts them at greater risk (28). The lack of habitat enrichment limits shelter-seeking behaviour in released European lobster, *Homarus gammarus* and those reared alone are slow to seek shelter (29), making them more vulnerable to predation (30). While training individuals alongside conspecifics can mitigate this (31), European lobsters are often reared individually to prevent agonistic interactions between individuals (32). With this vulnerability to predation in mind, it is important to maximise anti-predator defences prior to release.

European lobsters have attracted significant attention for restocking. Their populations collapsed due to overfishing during the 1970s (33) and several hatcheries have been established across Europe to help restock the natural population – for the benefit of both conservation and fisheries. Restocking is achieved in hatchery aquaria by rearing larvae through their planktonic and early benthic phases, keeping them in captivity during a time when, in the wild, they are most vulnerable to predation (33,34). Clawed lobster stocking programs use variable approaches, releasing lobsters from stage IV onwards into suitable sites to mature in their natural environment. Release strategies consider available shelter, as well as physical and oceanographic conditions required by European lobsters (35). Due to practical and logistical constraints, juvenile lobsters are often released during the day, when they will most easily be detected by visually guided predators. However, the visual components of the habitat (colour, pattern) have been neglected to date.

Juvenile European lobsters have seldom been observed in the wild, but those reared in hatchery aquaria show considerable individual variation in coloration with few resembling wild-caught adults (Fig 1). This presents a potential problem on release into the wild, as individuals are likely to be conspicuous to predators if poorly matched to their surroundings (1). Predation rates are highest in the first 24 hours following release (34), making this period a critical point in restocking programmes. Given that there is considerable variation in the colour of juvenile lobsters, knowing whether they can adapt to match the habitat, and whether aquarium colour can be altered to enhance habitat matching, should help hatcheries to enhance lobster anti-predator defences. If effective, such manipulations could have the potential to enhance survival on release. To date, no work has tested the capacity for this species to change colour and research on lobster coloration is limited to the influence of feed and genetics on pigmentation in the American lobster, *H. americanus* (36,37). Given the prevalence of background matching in crustaceans (3,38,39), it seems reasonable that juvenile lobsters could be capable of changing colour for the purposes of camouflage.

**Fig 1.**
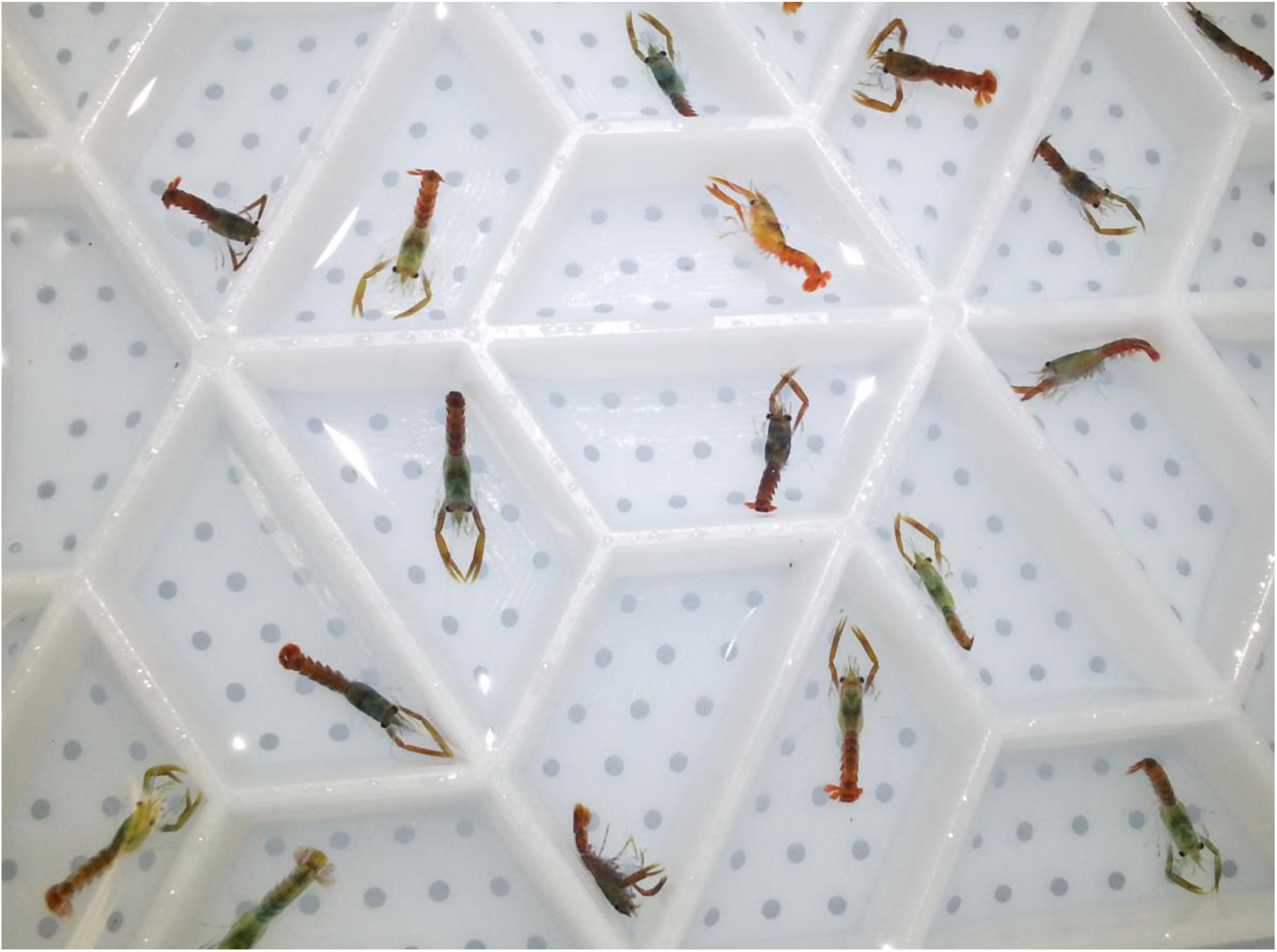
Variation in European lobster coloration (stage IV juveniles). The photo shows the variation in lobster coloration at the start of the experiment, following collection from the National Lobster Hatchery (NLH), Padstow, UK). Individuals here are housed within an Aquahive^®^ (Shellfish Hatchery Systems Ltd, Orkney, UK), a compartmentalised, stacked system used to rear juvenile European lobsters in captivity.

By placing hatchery-reared lobsters on artificial backgrounds and monitoring their coloration over time, we were able to test the ability of juvenile lobsters to change brightness in response to their surroundings. This study quantifies the capacity of lobsters to match their background using a model of fish vision (European pollack, *Pollachius pollachius*), allowing us to assess coloration from the predator’s perspective. Ultimately, this paper aims to assess whether altered hatchery practices can be used to improve lobster camouflage, and by implication survival, following release. We used a 35-day-long laboratory experiment to test the hypothesis that lobsters will change their coloration to better match their background over time, and that this will result in improvements in camouflage, when modelled to the visual systems of relevant predators.

## Materials and methods

### Husbandry

A total of 80 juvenile lobsters (all at stage IV, approximately 1 month old) were sourced from the National Lobster Hatchery (NLH) in Padstow (UK) and transported to the aquarium facility of the University of Exeter (Penryn Campus, UK). Lobsters were transported in an Aquahive^®^ tray (Fig 1), enclosed within a heavy-duty plastic bag containing seawater (32 +/− 2 ‰ salinity) and pure oxygen. This was secured within a 50 x 50 x 100 cm cool box to stabilise temperature during transport. Experimental glass tanks were set up to mimic hatchery rearing conditions as closely as possible. Water was supplied using uPVC push-fit pipe drilled with 1.5 mm diameter holes to allow aerated water to flow into each container. Before being recirculated, water was filtered (Classic 350 filter; Eheim GmbH & Co., Deizisau, Germany) and run through a heating system (DC300 Aquarium Chiller; D-D The Aquarium Solution Ltd., Ilford, UK) in order to maintain tank temperature (18-19 °C). Each tank was filled to 125 L with artificial seawater. Saltwater was made up to 32 +/− 2 ‰ salinity using Instant Ocean Salt (Instant Ocean, Blacksburg, Virginia) and dechlorinated water. Tanks were topped up with freshwater to compensate for evaporation during the course of the experiment. Tanks were prepared 48 hours before lobster arrival to allow them to reach a stable temperature.

Lobsters were housed individually (to prevent harm from aggressive interactions) in containers made from square uPVC gutter pipe (65 mm by 65 mm) cut to 60 mm lengths and covered with a mesh base to allow water through. All containers were fixed to corner braces and suspended within the tank above waterproof paper corresponding to the experimental treatment (a black or white background). Each juvenile was fed one formulated 1.5 mm diameter pellet daily. Pellets contained 133 mg/kg of the carotenoid Astaxanthin, known to affect coloration in a variety of crustaceans (40), including closely related species such as American lobster (37). All individuals were fed the same feed throughout the experiment to control for any influence of diet on lobster colour. Any uneaten food was removed the following day. Tanks were cleaned and half the water was changed twice weekly to limit any build up of bacteria and algae in the tanks. The light regime was set to 12 hours of light and 12 hours of darkness, with lights on from 07:30 to 19:30. Where handling was required, lobsters were pipetted between containers using a modified turkey baster, following the approach used by NLH. Containers were checked for moults twice daily on weekdays and once a day at weekends, confirming that individuals moulted during the experiment. Despite this, the precise number of moults undertaken by each individual is unknown owing to rapid consumption of the old exoskeleton following the moult. By the end of the experiment all individuals reached stage VI (determined by the white patterning at the edges of the carapace, known to develop during this stage).

### Experiment protocol

To determine the capacity of lobsters for background matching, juvenile lobsters were randomly assigned to either a black or a white compartment for a 2.5 or 5-week period. Individuals were photographed to determine their initial luminance (lightness as perceived by a particular predator, European pollack), and then photographed at various intervals (described below) to monitor changes in coloration over time. 80 individuals were used in the study; 40 of which were used to determine short (3 hours) and medium (2.5 weeks) term changes in coloration, and 40 of which were used to determine changes in luminance over the longer term (5 weeks). The allocation of individuals to each treatment is described in Table 1.

**Table 1:** **Experimental design.**

| Experiment | Aim | Black | White | Photography intervals |
| --- | --- | --- | --- | --- |
| Short-term | Quantify short-term change | n = 20 | n = 20 | 0 and 3 hours |
| Medium-term | Quantify plasticity & medium-term change | n = 20 | n = 20 | 0 and 2.5 weeks |
| Long-term | Monitor ontogenetic & longer-term change | n = 20 | n = 20 | 0, 2.5 and 5 weeks |

Treatments and recording intervals used in each experiment. The table outlines the number of juvenile European lobsters allocated to each treatment. The same individuals were used in both the short- and medium-term experiments.

Initial photographs of all 80 lobsters were taken 24 hours after introduction to the tank to prevent undue stress on individuals following transport. Further photos were taken after 2.5 weeks to quantify colour change over the medium-term. At this point, 40 individuals (20 of those on a white background and 20 of those on a black background) were transferred to the alternative treatment (black to white and *vice versa*) to assess reversible plasticity in colour change. The switched group were photographed after 3 hours and 2.5 weeks to determine the capacity for reversible changes in coloration over the short (3 hours) and medium (2.5 weeks) term. The group that remained on their original background were photographed after 2.5 and 5 weeks to assess longer-term changes in coloration. Lobsters were photographed in water within a 10 mm deep polytetrafluoroethylene (PTFE) chamber, under diffuse lighting conditions to minimise stress during data collection. Lobsters that moulted on the same day that photography was scheduled were not photographed to prevent damage to the new exoskeleton during handling. The effect of moulting on coloration was not quantified given that multiple moults occurred between each photography interval and it was not possible to record every moulting event (as described above). Consequently, it was not possible to distinguish between colour change within moults (e.g. due to pigment dispersal) and between moults, both of which are responsible for colour change in crustaceans (3,39,41).

### Quantifying colour change

Both colour change and camouflage was quantified with respect to predator vision (42–44). Photos were taken using a Nikon D7000 SLR, fitted with a 60 mm quartz lens (Coastal Optics). A 400-700 nm, Baader Venus U filter was fixed in front of the lens, allowing wavelengths visible to European pollack to be recorded. All photos were taken under simulated daylight conditions, achieved using an arc lamp (Ventronic) equipped with a daylight 65 bulb. All lobsters were photographed against a white background in a clear PTFE chamber. A translucent white PTFE diffuser was placed between the photography chamber and the light source to ensure lighting was even. Two grey standards (7% and 93% reflectance) were used in every photo to account for any variation in illumination over time, following methods developed by Troscianko and Stevens (44). The camera white balance was set to manual and the aperture was kept constant between photos. All photographs were taken during daylight hours to account for any potential coloration with day-night cycles, as has been observed in other crustaceans (45–47).

### Image analysis

To establish how lobsters would be perceived by predators, the images were mapped to fish vision (European pollack) using an established polynomial mapping technique (44), which yields predicted cone catch data that are highly accurate compared to data obtained *via* reflectance spectrometry (44,48,49). Images were analysed in ImageJ (National Institute of Health, NIH) using the Multispectral Image Calibration and Analysis Toolbox (44). The longwave channel was used to calculate luminance (lightness according to a specific visual system) as potentially perceived by a dichromatic predatory fish, using spectral sensitivity data from European pollack (50). Average luminance was calculated for each lobster by selecting a rectangular region of interest (ROI) that covered as much of the carapace as possible while excluding the white patterning observed at the edges of the cephalothorax. This patterning develops with age and was excluded from the ROI so that reversible plasticity in coloration could be quantified.

To assess the lobsters’ level of camouflage, a widely-implemented predator discrimination model, based on receptor-noise limited discrimination, was used (51,52). This calculates Just Noticeable Differences (JNDs) between one object (the lobster) and another (the background) to predict how easily the two can be distinguished, according to a particular visual system (here, European pollack). Luminance JNDs were calculated using a modified (log) version of the Vorobyev and Osorio model for luminance discrimination (51,52), using a Weber fraction of 0.05, which is thought to be appropriate for many fish (53). This approach has been used to quantify camouflage in several studies of animal coloration, including the perception of crustaceans by European pollack (54,55).

### Statistical analysis

All statistical analyses were carried out using R version 3.31 (56). Linear mixed effects models were used to assess the effect of time, background colour (black, white) and their interaction on luminance, and to assess the effect of time on camouflage (expressed in JNDs (51)). Lobster ID was included as a random effect in all models in order to account for any potential temporal autocorrelation. For all models, the following approach was used: linear mixed models were fitted by restricted maximum likelihood (REML), with Kenward-Roger approximations to degrees of freedom using the lme4 package (57). The minimum adequate model was determined by successively removing non-significant terms, starting with the highest order terms in the model. Analysis of variance (ANOVA) was used to determine which model best explained the variation in the response. Assumptions of normality were supported for all datasets, which were checked through visual inspections of quantile-quantile plots, residual distributions and residual vs fitted values plots. In addition, Welch two sample t-tests were used to evaluate differences in mean luminance change between lobsters allocated to a black background and those allocated to a white one.

## Results

### Short-term change

Juvenile lobsters show no significant response to their background in the short-term; the only significant term was the intercept (GLM: t = 20.46, d.f. = 42, p < 0.001), resulting in a null model (see S1 Table for model output). After 3 hours of exposure (S1 Table), there was no significant interaction between time and background (ANOVA: Chi-sq = 3.07, d.f. = 1, p = 0.080), no significant effect of background on coloration (ANOVA: Chi-sq = 0.02, d.f. = 1, p = 0.889) and no significant effect of time (ANOVA: Chi-sq = 2.40, d.f. = 1, p = 0.121), resulting in a null model.

### Medium-term change

During the first 2.5 weeks in the laboratory, lobsters were observed to darken over time (GLM: t = 16.53, d.f. = 39, p <0.001), (Fig 2A), with individuals on a darker background as dark as those on a light one, on average (t-test: t = 1.06, d.f. = 37, p = 0.298), (Fig 2B). However, neither the interaction between time and background (ANOVA: Chi-sq = 1.16, d.f. = 1, p = 0.282) nor background as an independent variable (ANOVA: Chi-sq = 1.32, d.f. = 1, p = 0.250) had a significant effect on lobster coloration. Model parameters are detailed in Table 2.

**Fig 2.**
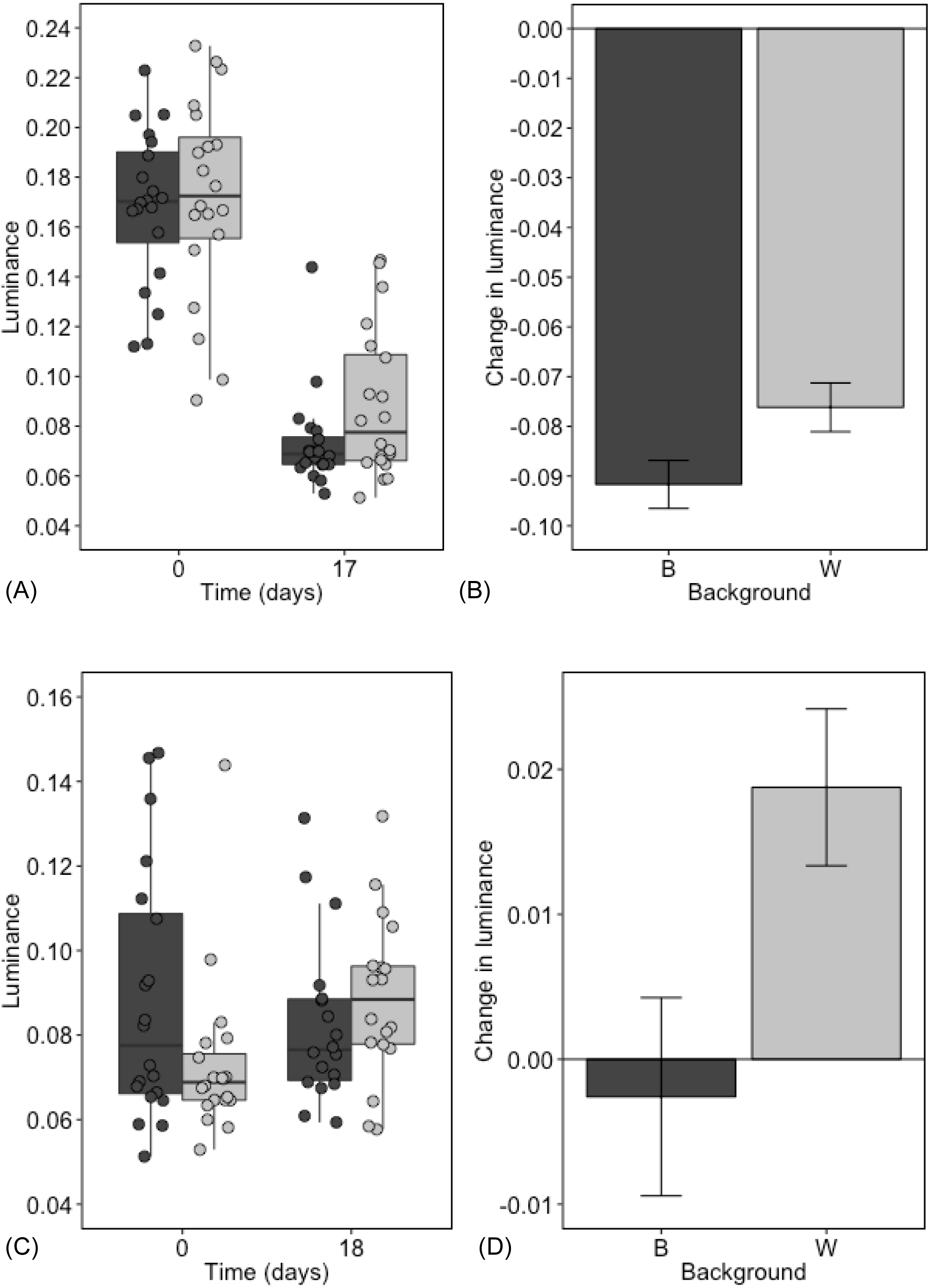
Change in juvenile lobster luminance in response to black and white backgrounds over the medium term. Dark and light grey points, boxes and bars correspond to lobsters on a black and white background, respectively. Panels (A) and (B) show the initial luminance change observed in lobsters placed on a black or white background; panels (C) and (D) show the plastic change following transfer to the second background treatment (those initially on black were transferred to white and *vice versa*). Panels (A) and (C) show the variation in luminance across the experimental population, where central lines are medians, boxes are interquartile ranges and whiskers are 95% quartiles. Panels (B) and (D) show the mean luminance change observed following exposure to the background treatments, together with standard errors. Luminance is presented according to European pollack vision.

**Table 2:**
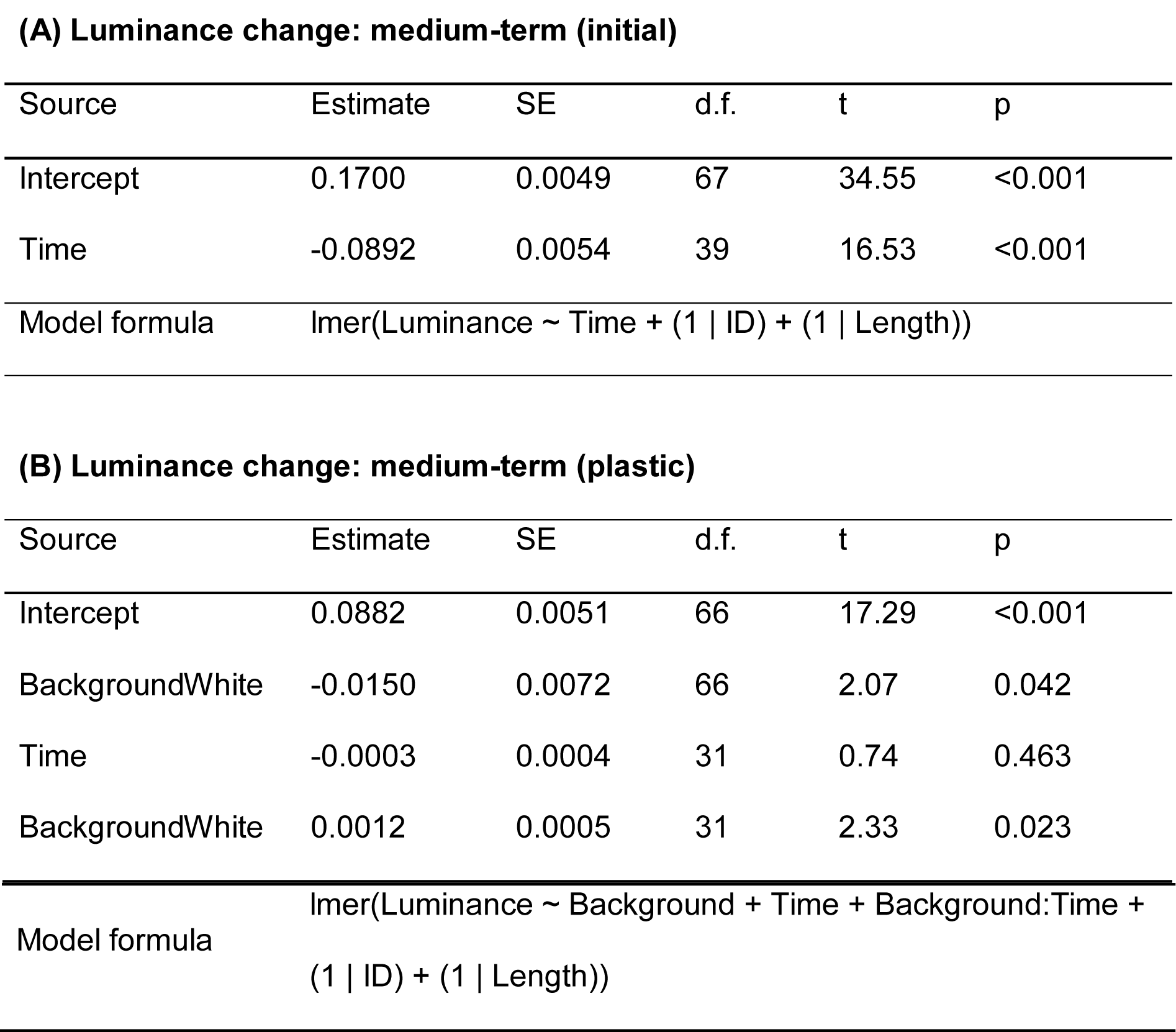
**Parameter estimates from the minimum adequate model describing the change in juvenile lobster luminance in response to black and white backgrounds over the medium term.**

**(A) Luminance change: medium-term (initial)**
| Source | Estimate | SE | d.f. | t | p |
| --- | --- | --- | --- | --- | --- |
| Intercept | 0.1700 | 0.0049 | 67 | 34.55 | <0.001 |
| Time | -0.0892 | 0.0054 | 39 | 16.53 | <0.001 |
| Model formula | lmer(Luminance ~ Time + (1 \| ID) + (1 \| Length)) |  |  |  |  |

**(B) Luminance change: medium-term (plastic)**
| Source | Estimate | SE | d.f. | t | p |
| --- | --- | --- | --- | --- | --- |
| Intercept | 0.0882 | 0.0051 | 66 | 17.29 | <0.001 |
| BackgroundWhite | -0.0150 | 0.0072 | 66 | 2.07 | 0.042 |
| Time | -0.0003 | 0.0004 | 31 | 0.74 | 0.463 |
| BackgroundWhite | 0.0012 | 0.0005 | 31 | 2.33 | 0.023 |

|  |  |
| --- | --- |
| Model formula | $\text{lmer}(\text{Luminance} \sim \text{Background} + \text{Time} + \text{Background}:\text{Time} + (1 \vert \text{ID}) + (1 \vert \text{Length}))$ |

In all cases luminance is expressed according to European pollack vision. Initial luminance change (A) describes the change in luminance when placed on the first background (black or white). Plastic luminance change (B) describes the change in luminance when placed on the second background (individuals on a black background were transferred to white and *vice versa*). Linear mixed models were fitted by restricted maximum likelihood (REML) using the lme4 package (57). The Kenward-Roger approximation for degrees of freedom was used to determine p-values. Lobster ID and length were included as random effects.

Despite initially darkening over time (Fig 2A,B), lobsters showed some capacity to change luminance in response to their surroundings when switched to the alternative background type for a further 2.5 weeks. The interaction between time and background had a significant effect on luminance (GLM: t = 2.33, d.f. = 31, p = 0.026), (Fig 2C). Lobsters on a black background darkened whereas those on a light background became lighter, on average (t-test: t = 2.45, d.f. = 32, p = 0.020), (Fig 2D). The full model containing the interaction between time and background was significantly better than simpler alternatives (ANOVA: Chi-sq = 5.27, d.f. = 1, p = 0.022); model parameters are detailed in Table 2B.

The initial darkening observed in both treatments (Fig 2A,B) corresponds to a significant decrease in camouflage over time for lobsters on a white background (GLM: t = 10.14, d.f. = 19, p < 0.001), (Fig 3A) and a significant increase in camouflage for those initially placed on a black background (GLM: t = 13.65, d.f. = 19, p < 0.001), (Fig 3B). In both cases, the full model containing time, background and their interaction was a significantly better explainer for the variation in camouflage than simpler alternatives (ANOVA_white_: Chi-sq = 40.68, d.f. = 1, p = <0.001; ANOVA_black_: Chi-sq = 62.55, d.f. = 1, p = <0.001).

**Fig 3.**
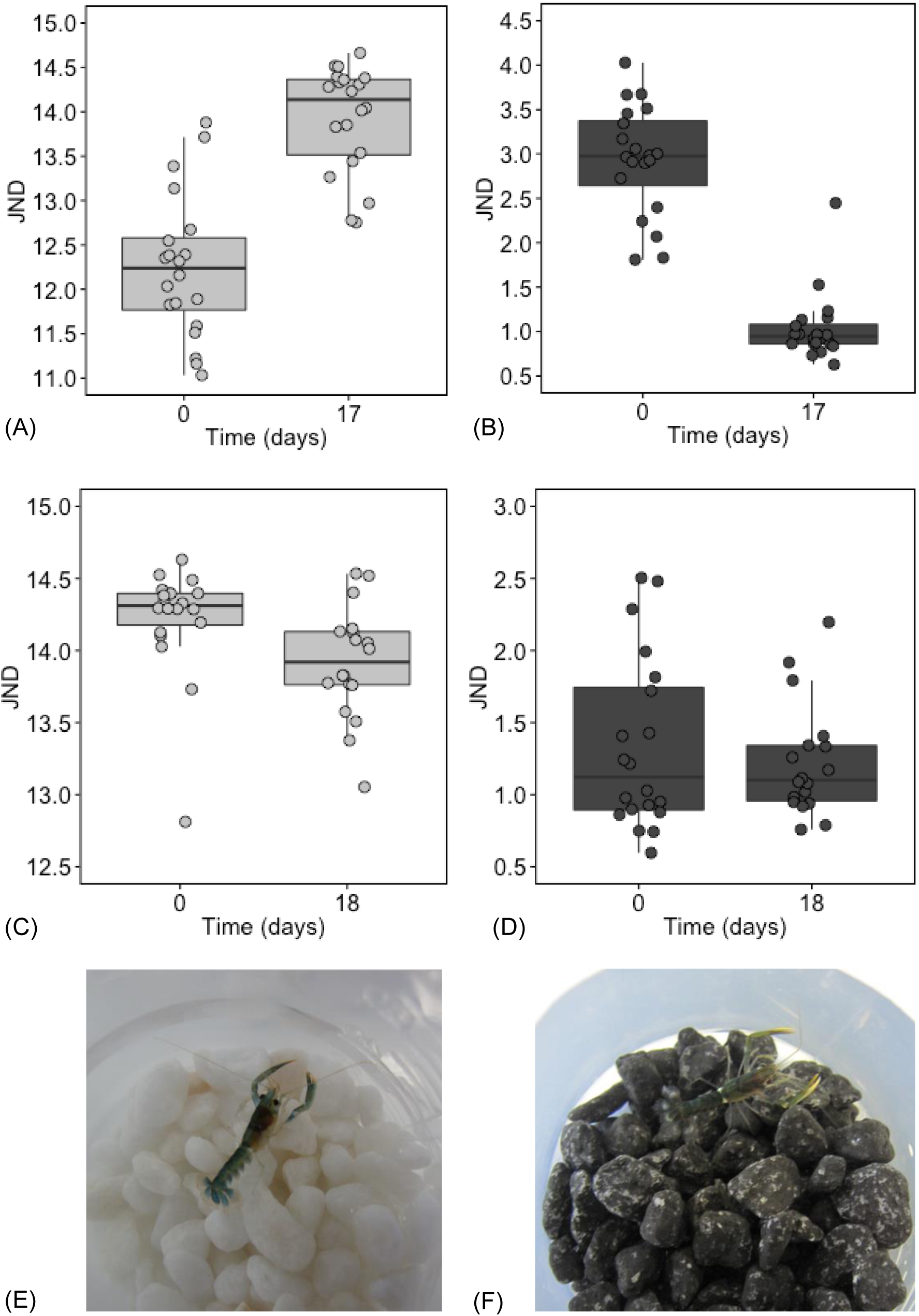
Change in juvenile lobster camouflage against black and white backgrounds over the medium term. Panels on the left correspond to individuals allocated to a white background (light grey points and boxes) and those on the right correspond to individuals allocated to a black background (dark grey points and boxes). Camouflage is presented according to European pollack vision. The change in detectability is quantified using Just Noticeable Differences (JNDs), (51) and is shown for both initial changes in coloration (A, B) and for individuals placed on the alternative background (C, D). A decline in JND corresponds to a predicted decrease in detectability according to predator vision (i.e. an increase in camouflage). JNDs of 1 or below correspond to objects (a lobster, its background) that cannot be distinguished from each other (52). Central lines are medians, boxes are interquartile ranges and whiskers are 95% quartiles. Both (E) and (F) are example individuals, showing level of lightness/darkness attained after 18 days on their respective backgrounds.

Plastic changes in coloration (Fig 2C,D) corresponded to a small but significant increase in camouflage over time for those on a white background (GLM: t = 2.45, d.f. = 36, p = 0.020), (Fig 3C). However, no significant change in camouflage was seen in individuals allocated to a black background (GLM: t = 13.38, d.f. = 18, p < 0.001), (Fig 3D). In this instance, the interaction between background and time was not significant (ANOVA: Chi-sq = 0.46, d.f. = 1, p = 0.499), likely because individuals were already quite dark (Fig 3). Individuals allocated to a black background were already well matched to their surroundings at the start of the plastic trial (JND close to one, Fig 3D). Model parameters are detailed in Table 3 (see S2 Table for mean JNDs).

**Table 3:**
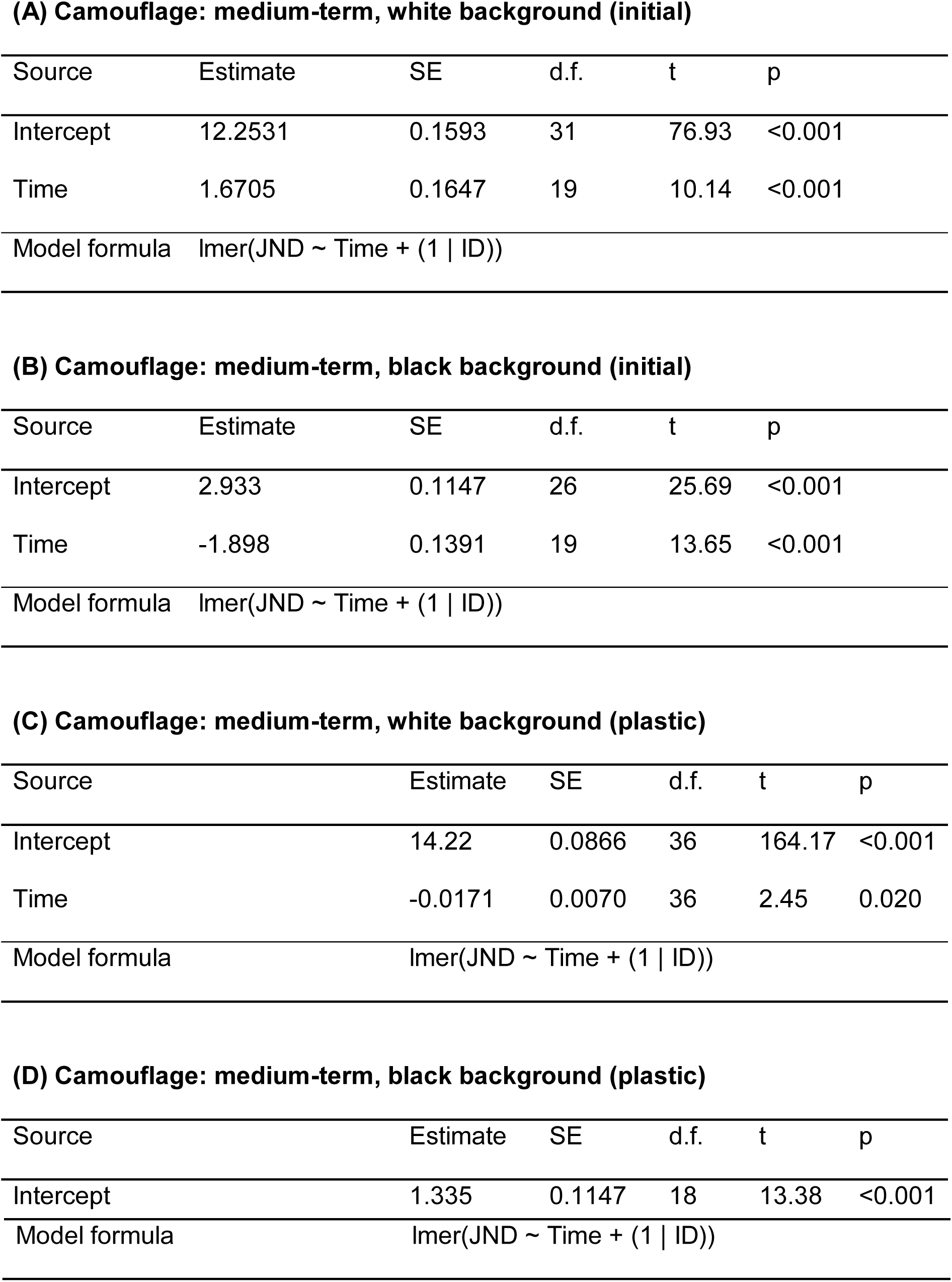
**Parameter estimates from the minimum adequate models describing the change in juvenile lobster camouflage against black and white backgrounds over the medium term.**

Initial change (A, B) describes the change in camouflage when placed on the first background (black or white). Plastic change (C, D) describes the change in camouflage when placed on the second background (individuals on a black background were transferred to white and *vice versa*). Camouflage is expressed in Just Noticeable Differences (JNDs), a measure of discriminability according to predator (European pollack) vision. Parameter estimates for individuals allocated to a white background (A, C) and black background (B, D) are shown. Linear mixed models were fitted by restricted maximum likelihood (REML) using the lme4 package (57). The Kenward-Roger approximation for degrees of freedom was used to determine p-values. Lobster ID was included as a random effect.

### Long-term change

Lobsters that were not presented with a new background darkened significantly throughout their time in the laboratory (GLM: t = 10.35, d.f. = 107, p < 0.001), (Fig 4). However, neither the interaction between time and background (ANOVA: Chi-sq = 0.04, d.f. = 1, p = 0.834) nor background as an independent variable (ANOVA: Chi-sq = 2.43, d.f. = 1, p = 0.119) had a significant effect on luminance. Model parameters are detailed in Table 4. Lobsters darkened by the same extent over a 35-day period, regardless of their background (t-test: t = 0.06, d.f. = 31, p = 0.956), (Fig 4 insert). This darkening resulted in an increase in detectability for those on a light background (GLM: t = 8.38, d.f. = 34, p <0.001), (Fig 5A) and decrease in detectability for those on a black one (GLM: t = 6.34, d.f. = 56, p < 0.001), (Fig 5B). Model parameters are summarised in Table 5.

**Fig 4.**
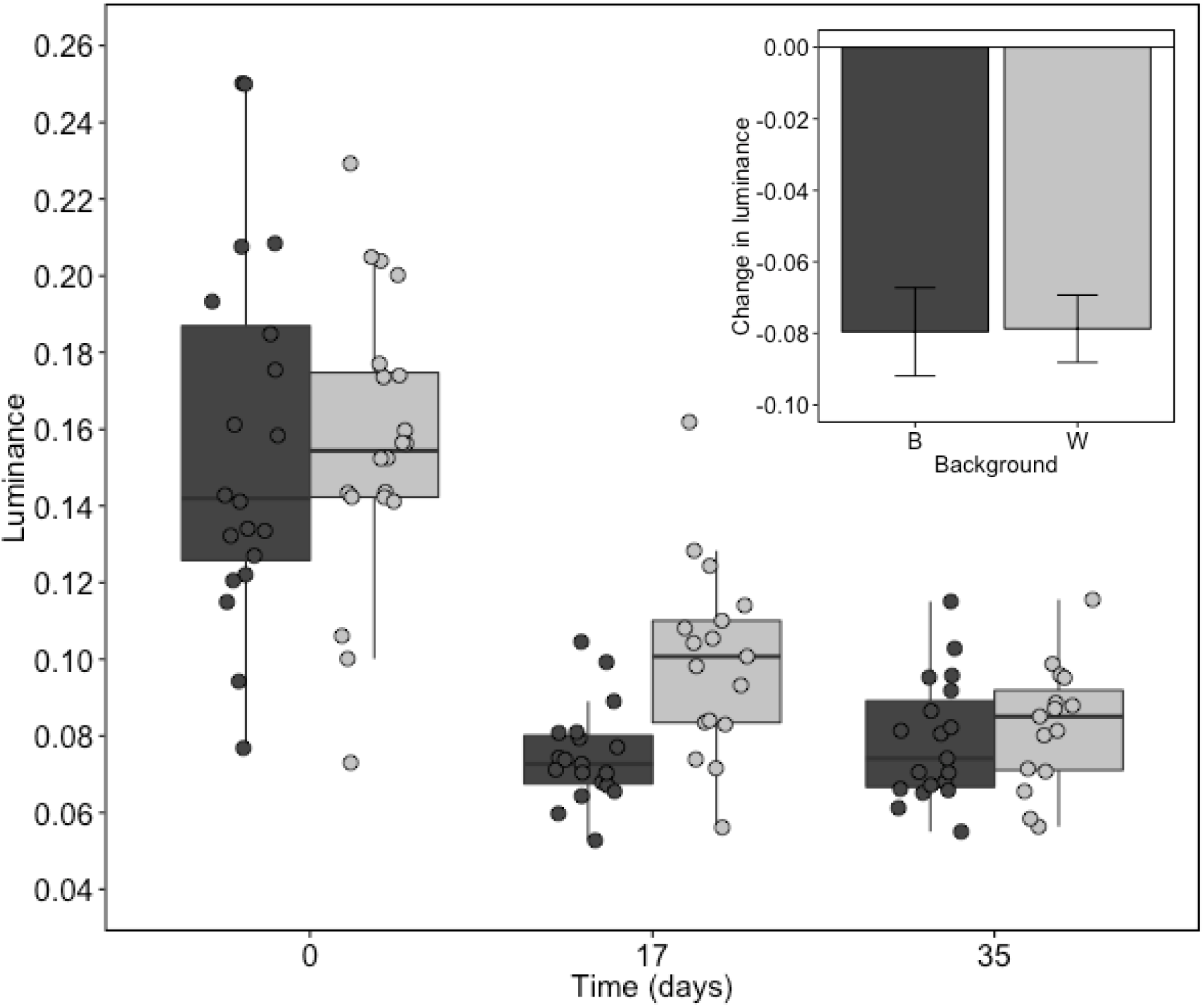
Change in juvenile lobster luminance in response to black and white backgrounds over the long term. Dark and light grey points, boxes and bars correspond to lobsters on a black and white background, respectively. The boxplot shows the variation across the experimental population at each time point, where central lines are medians, boxes are interquartile ranges and whiskers are 95% quartiles. The insert shows the mean change in lobster luminance observed for each background treatment over the 5-week experimental period (luminance on day 35 – luminance on day 0). Error bars in insert show standard errors. Luminance is presented according to European pollack vision.

**Table 4:**
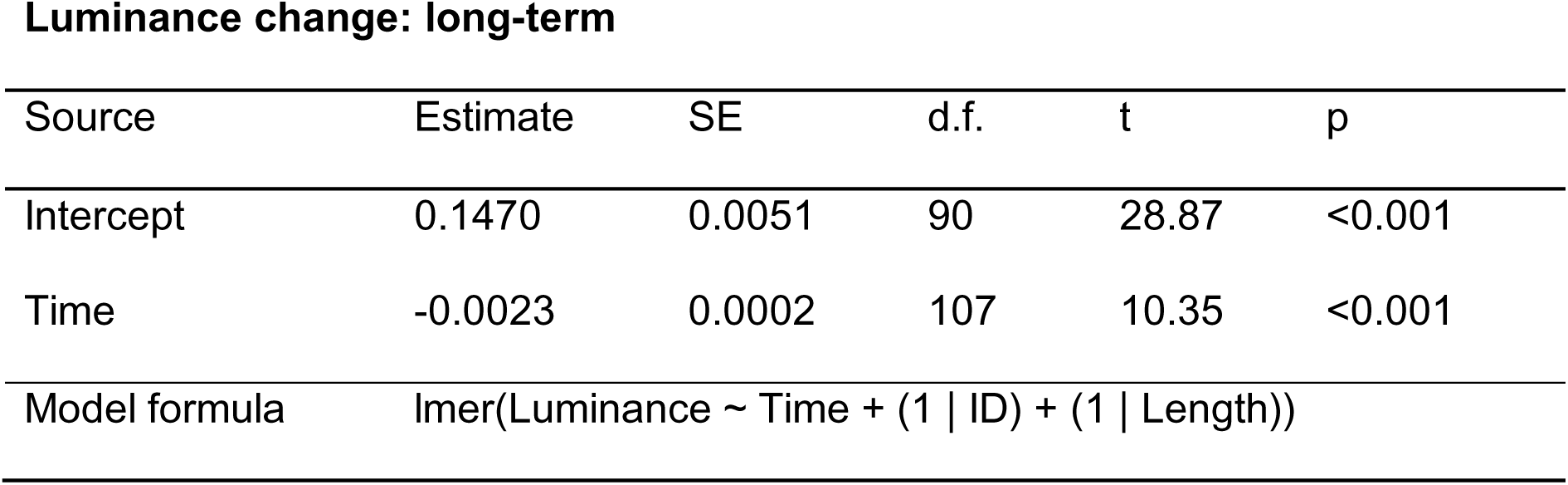
**Parameter estimates from the minimum adequate model describing the change in juvenile lobster luminance in response to black and white backgrounds over the long term.**

| Source | Estimate | SE | d.f. | t | p |
| --- | --- | --- | --- | --- | --- |
| Intercept | 0.1470 | 0.0051 | 90 | 28.87 | <0.001 |
| Time | -0.0023 | 0.0002 | 107 | 10.35 | <0.001 |
| Model formula | lmer(Luminance ~ Time + (1 \| ID) + (1 \| Length)) |  |  |  |  |

**Fig 5.**
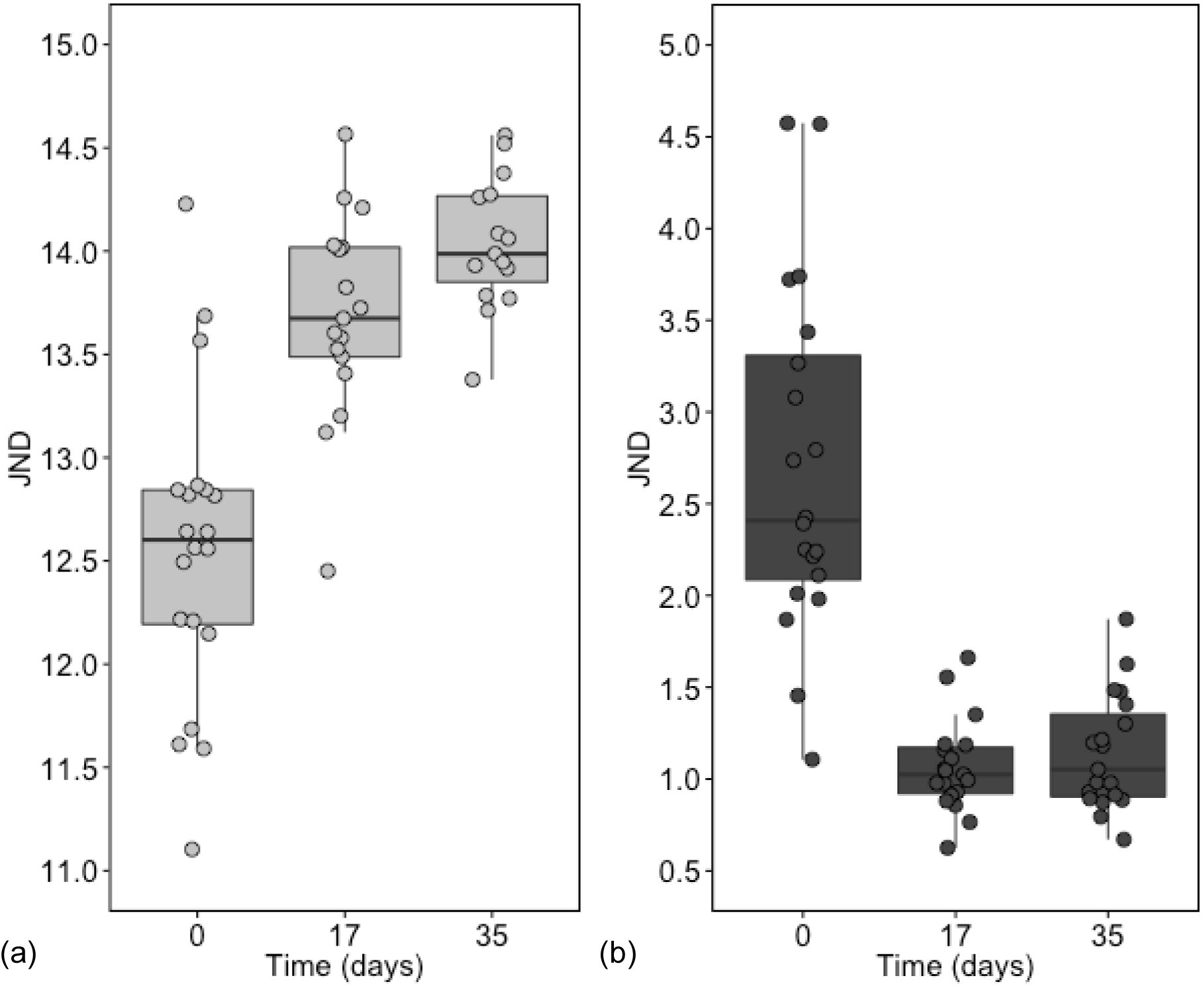
Change in juvenile lobster camouflage against black and white backgrounds over the long term. Camouflage is presented according to European pollack vision. The change in detectability according to Just Noticeable Differences (JNDs) (51) is shown for (A) juvenile lobsters placed on a white background (light grey points and boxes) and (B) for those on black (dark grey points and boxes). Central lines are medians, boxes are interquartile ranges and whiskers are 95% quartiles. Note that a decline in JND corresponds to a decrease in predicted detectability according to predator vision (i.e. an increase in camouflage) and that JNDs of 1 or below correspond to objects (here a lobster and its background) that cannot be distinguished from each other (52).

**Table 5:** **Parameter estimates from the minimum adequate models describing the change in juvenile lobster camouflage against black and white backgrounds over the long term.**

**(A) Camouflage: long-term, white background**
| Source | Estimate | SE | d.f. | t | p |
| --- | --- | --- | --- | --- | --- |
| Intercept | 12.68 | 0.1265 | 41 | 100.3 | <0.001 |
| Time | 0.0432 | 0.0052 | 34 | 8.38 | <0.001 |
| Model formula | lmer(JND ~ Time + (1 \| ID)) |  |  |  |  |

**(B) Camouflage: long-term, black background**
| Source | Estimate | SE | d.f. | t | p |
| --- | --- | --- | --- | --- | --- |
| Intercept | 2.410 | 0.1489 | 56 | 16.19 | <0.001 |
| Time | -0.0444 | 0.0067 | 56 | 6.64 | <0.001 |
| Model formula | lmer(JND ~ Time + (1 \| ID)) |  |  |  |  |

Linear mixed models were fitted by restricted maximum likelihood (REML) using the lme4 package (57). The Kenward-Roger approximation for degrees of freedom was used to determine p-values. Lobster ID and length were included as random effects.

Tables (A) and (B) show changes in camouflage for individuals allocated to a white background and black background, respectively. Camouflage is expressed in Just Noticeable Differences (JNDs), a measure of discriminability according to predator (European pollack) vision. Linear mixed models were fitted by restricted maximum likelihood (REML) using the lme4 package (57). The Kenward-Roger approximation for degrees of freedom was used to determine p-values. Lobster ID was included as a random effect.

## Discussion

Juvenile European lobsters show no significant change in coloration in response to their background in the short-term (3 hours). The absence of rapid camouflage in this species may contribute to the high predation mortality observed in the first 24 hours following release into the wild (34). Given that individuals do not change their coloration rapidly, if they are reared on substrates that are not representative of the natural environment, they may stand out against wild substrates, making them an easy target for predators. Luminance (lightness as perceived by a particular predator) is a strong contributor to visual detectability. When transferred to a new substrate, individuals became lighter (increasing in luminance) on a white background and darker on a black one over 2-3 weeks (Figs 2C,D and 3). This medium-term response to their substrate shows that juvenile European lobsters are capable of some degree of background matching and, over time, have the potential to become increasingly difficult to discern from their surroundings (Fig 3). However, some of the changes in discriminability are small (less than 1 JND for those on a white background), so may not always be perceptible to predator visual systems. More effective changes in camouflage may be achieved or through the use of more natural substrates during rearing or with earlier exposure to the background. While black and white features like pebbles occur in the environment, the backgrounds used here were artificial and do not represent the wide range of substrates that would be encountered in nature. It is entirely possible that when faced with more naturalistic substrates (resembling naturally occurring rock, sediment or algae) that lobsters may show a greater degree of plasticity and matching. Greater background matching could also occur if individuals are exposed to different substrates as soon as they settle from the plankton, when, in accordance with their life history, there would be the greatest need for habitat matching. The gradual changes in response to the background observed here (Fig 2C,D) may correspond to improvements in camouflage (Fig 3) and, potentially, survival (17,18), but because colour change in European lobster is relatively slow (Figs 2 and 3), any benefit would require pre-conditioning individuals for the habitat they are released into. Post-release survival will also depend on the habitat and predators present at the release site. Note that habitat matching is one of many factors that should be considered when determining how to maximise chances of lobster survival on release and that other physical factors, such as nutritional state and fitness should also be optimised prior to release.

In both the medium- and long-term experiment, individuals darken initially, with most of that darkening occurring within the first 2.5 weeks (Figs 2 and 4). Individuals that remained on the same background became darker over a longer time period (5 weeks), regardless of their background treatment (Fig 5). Given that darkening occurs over the longer-term, it likely reflects an ontogenetic change in coloration (58). Ontogenetic changes in coloration may reflect a number of drivers, including relaxed selection pressure on coloration (e.g. fewer predators and less need for camouflage), movement into new habitats with age, or changes in camouflage strategy (16,58–60). In many animals, predation risk changes with age as individuals gain a size refuge from predation. The risk is greatest for younger, smaller individuals and lessens as individuals grow and develop weapons, (such as chelae) for defence, or a more robust body morphology (16,61). As such, there may be stronger selective pressure for effective anti-predator defences, like camouflage, in early life stages (3). Ontogenetic changes in coloration can also coincide with changes in habitat use throughout development (16,62), as different colours and patterns may provide better camouflage in different habitats. Lobsters are initially pelagic in their early life stages, but these early stages are poorly understood. Pelagic juveniles may remain in the open ocean, or congregate close to shore and vary their position in the water column with age. In the brightly light surface oceans it pays to be lightly coloured or even translucent. Pelagic juveniles in coastal waters would also benefit from being pale or translucent, but may be more varied in colour depending on the overall appearance of the habitat (the dominant algae or bedrock, for example). Regardless of the habitat experienced by planktonic juveniles, when they reach the deeper, darker seabed they will need to be correspondingly dark, so it stands to reason that settlement could act as a trigger for darkening. Long-term darkening is slightly, and temporarily, offset by the plastic response to their background in the medium-term, with those on a white background becoming less dark than those on black (Fig 4). This indicates that rearing benthic stages in white containers in the hatchery may marginally impede individuals from darkening as they would in a more natural (darker) habitat. However, the impact of this on detectability depends on the time spent in the rearing container, as over longer periods all lobsters were seen to darken over time (Fig 4).

Planktonic lobster larvae can disperse over a large area (63), with single populations extending up to 230 kilometres (64). Given this extensive larval dispersal, plasticity in coloration (in order to match the local environment after settling) has clear advantages. This study highlights the potential for juvenile lobsters to change coloration (lightness / darkness) to better match their surroundings, but further work is needed to quantify their capacity for adaptive camouflage in response to natural backgrounds. For example, their ability to change colour as well as luminance, which could afford them protection from predators in a variable environment (54,65) and across the range of habitats into which they might settle. Furthermore, the ability to fine-tune their lightness to match a specific habitat may have added benefits for individuals as they age, given the site-fidelity exhibited by European lobsters (66,67). If reared in an environment that resembles their release site, rather than a bright white background, individuals may be more likely to evade predator detection when released into the wild. Ecological drivers of colour change have the potential to provide nature-based solutions for stocking, but further work is needed to ascertain whether colour change in this species corresponds to an improvement in survival. We encourage researchers to explore the drivers of changes in coloration in this species and the potential for changes in hue, as well as luminance, in juvenile lobsters. Ontogenetic changes are likely to override such plasticity longer term, as adult European lobsters are much less variable than juveniles. Such shifts from plastic camouflage to ontogenetic changes in coloration have been observed in other marine crustaceans (16,68). These ontogenetic shifts in coloration correspond to improved camouflage across the range of environments experienced by adults (68). Consequently, the timing of rearing individuals on different backgrounds relative to the timing of release needs careful thought in order to maximise any applied benefits of camouflage for stocking.

There is potential for camouflage to be further enhanced through diet modification. Significant work has explored the optimal diet for rearing fast-growing, healthy juveniles in hatchery conditions (69,70), but the impact of diet on coloration in this species is largely unknown. Experiments on American lobster have revealed variation in coloration depending on the amount of carotenoids in the diet, with lobsters fed on a low carotenoid diet appearing blue, and those fed on a diet high in carotenoids being redder in colour (37). However, larval lobsters do not show any changes in response to carotenoid-rich diets (71), suggesting that carotenoid availability only influences coloration beyond the larval stage. With this in mind, diet may play an important role in ontogenetic changes in appearance. The amount dietary carotenoids is also known to affect coloration in other species, including giant tiger prawns, with more carotenoids resulting in darker coloration (72). Many species obtain pigment through ingestion in order to better match their background including caterpillars (73), spiders (74) and, potentially, prawns (3). Diet modification, and the provision of foodstuffs resembling those naturally found at release site, may further enhance the camouflage and hence survival of hatchery-reared juveniles.

The findings presented here may also have implications for shellfish product coloration, particularly given recent advances in the potential for aquaculture of this species (75). Individuals with darker, more striking coloration often attract a higher price (76). If the plasticity in coloration seen here exists in later developmental stages, then responses to background colour and brightness could be harnessed to produce higher value shellfish for market. While some individuals in our study converged on the same brightness in the longer-term (Fig 4), on other backgrounds and with differences in diet more plasticity may be possible. These considerations apply not only to lobster, but also other crustacean aquaculture, as many commercial species are capable of colour change, including giant tiger prawns, *Penaeus monodon*, and Pacific white shrimp, *Litopenaeus vannamei*, and are more valuable when exhibiting a particular colour (40,72,76,77). Understanding and applying camouflage offers potential advantages to stocking and aquaculture programs, as individuals reared in environments that appear natural may be more natural in colour. Natural coloration will likely afford individuals reared for conservation and stock enhancement with protection from predators on release. Releasing individuals with effective anti-predator defences is vital to ensure the success of stocking programs. Similarly, as either more natural or darker coloration can enhance product value, aquaculturists stand to benefit from modifying rearing environments to promote such traits, particularly as the economic viability of aquaculture ventures depends on product value (72,76).

This study is the first to demonstrate the capacity of early benthic phase lobsters to change brightness in response to their background, and show that these changes can affect detectability according to models of predator vision. These findings have direct relevance to restocking programmes, which aim to maximise the fitness of captive-reared lobsters on release. Further work should test the capacity of early benthic phase larvae to match the colour of ecologically relevant substrates, such as sand, mud, cobbles and seaweed (78–80). Understanding the implications of hue as well as brightness on lobster coloration will greatly inform our understanding of phenotypic plasticity in this species and determine the extent to which rearing environments can be modified to increase anti-predator defences in the natural environment. With refinement of these approaches, there is potential for colour change to be harnessed to improve lobster camouflage prior to release. Given the high predation rate experienced immediately following release, such measures may potentially improve the survivorship of released juveniles. However, precise timings should consider when plasticity is highest during development. Further work is needed to determine whether conservation projects and aquaculture programmes may reap such benefits through careful selection of artificial substrates and altering the colour of rearing environments to enhance camouflage.

## Acknowledgements

Thanks to National Lobster Hatchery staff for their support throughout this project, particular thanks to Charlie Ellis for his help initiating the work. Thanks also to Steve Votier and Mark Briffa for their insightful comments on an earlier version of this manuscript.

