## Supplementary tables for "Using camouflage for conservation: colour change in juvenile European lobster"

*Corresponding author

**S1 Table. Parameter estimates from the minimum adequate model describing the change in juvenile lobster luminance in response to black and white backgrounds over the short term.**

**Luminance change: short-term**

| Source | Estimate | SE | d.f. | t | p |
|---|---|---|---|---|---|
| Intercept | 0.0809 | 0.0040 | 42 | 20.46 | <0.001 |
| Model formula | lmer(JND ~ Time + (1 \| ID)) | | | | |

Short-term (3-hour) changes in juvenile lobster luminance (lightness according to European pollack vision) for individuals allocated to a black or white background. Linear mixed models were fitted by restricted maximum likelihood (REML) using the lme4 package (1). The Kenward-Roger approximation for degrees of freedom was used to determine p-values. Lobster ID and length were included as random effects.

**S2 Table. Mean change in lobster camouflage according to pollack vision observed over the medium term.**

**(A) Mean JND: white background (initial)**

| Time (days) | N | Mean JND | SD | SE |
|---|---|---|---|---|
| 0 | 20 | 12.25 | 0.81 | 0.18 |
| 17 | 20 | 13.92 | 0.60 | 0.13 |

**(B) Mean JND: black background (initial)**

| Time (days) | N | Mean JND | SD | SE |
|---|---|---|---|---|
| 0 | 20 | 2.93 | 0.61 | 0.14 |
| 17 | 20 | 1.04 | 0.39 | 0.09 |

**(C) Mean JND: white background (plastic)**

| Time (days) | N | Mean JND | SD | SE |
|---|---|---|---|---|
| 0 | 20 | 14.22 | 0.39 | 0.09 |
| 18 | 18 | 13.91 | 0.39 | 0.09 |

**(D) Mean JND: black background (plastic)**

| Time (days) | N | Mean JND | SD | SE |
|---|---|---|---|---|
| 0 | 20 | 1.33 | 0.60 | 0.13 |
| 18 | 18 | 1.23 | 0.39 | 0.09 |

Camouflage is expressed in Just Noticeable Differences (JNDs) (2), a measure of discrimination according to predator (pollack) vision, where a decrease in JND corresponds to an increase in camouflage. Both (A) and (B) show the mean JND for individuals placed on their first background: (A) white, (B) black; and (C) and (D) show the mean JND for individuals placed on the alternative background treatment: (C) white, (D) black.
